# Fecal Virome Transplantation (FVT) from healthy humans remodels the gut bacteriome and virome and reduces metabolic syndrome in mice

**DOI:** 10.1101/2024.02.17.580809

**Authors:** Melany J Cervantes-Echeverría, Marco Antonio Jimenez-Rico, Rubiceli Manzo, Abigail Hernández-Reyna, Fernanda Cornejo-Granados, Filiberto Sánchez-López, Jonathan Salazar-León, Gustavo Pedraza-Alva, Leonor Perez-Martinez, Adrian Ochoa Leyva

## Abstract

The gut microbiome regulates host processes, including metabolism, and comprises viruses, archaea, fungi, protists, and bacteria. However, previous studies have primarily focused on the bacterial components of the microbiome. This study conducted cross-infection experiments by fecal virome transplantation (FVT) to examine the impact of virus-like particles (VLPs) from normal-weight individuals on bacterial and virome diversity and metabolic changes in mice. We found that FVT significantly restored glucose tolerance in mice, even though they were maintained on a high-fat diet (HFD). Shotgun metagenomics and 16s profiling confirmed that FVT shifted gut bacterial and viral richness and diversity, resulting in several differentially abundant bacteria and phages. The abundance of Akkermansia muciniphila decreased significantly after FVT, remained low until the end of the experiment (week 17), and correlated with a decreased glucose tolerance. On the other hand, there was a significant increase in Allobaculum and Coprococcus at weeks 10 and 17 post-FVT. The abundance of Allobaculum correlated with an increased glucose tolerance, indicating a beneficial hypoglycemic effect on the mice inoculated with VLPs. We also found an increased virome diversity at week 17, while richness was similar to the one before FVT. The study provides a proof of concept for the efficacy of FVT from human donors to mice, improving glucose tolerance by altering the gut microbiota and virome despite maintaining a HFD. Overall, our results suggest that trans-kingdom interactions between the virome and bacteriome from humans could be translated to mice as a study model.

## Introduction

Obesity is a leading global nutritional and public health problem caused by excessive body fat accumulation due to an imbalance between energy intake and expenditure [1, 2]. It can lead to serious health issues, including metabolic syndrome (MetS), a condition where an individual has three or more risk factors for health, such as hyperglycemia, hypertriglyceridemia, low high-density lipoprotein or HDL cholesterol levels, high blood pressure, and abdominal obesity [3, 4].

Most studies have focused on the gut-associated bacterial component associated with obesity and MetS [5]. However, recent studies suggest that changes in the gut virome also play an essential role in these conditions [6, 7]. Understanding the interrelationship between gut bacteria and virome associated with obesity and MetS can help us address this disease and related health risks.

The human gut viral community, the virome, mainly comprises bacterial viruses called bacteriophages or phages [6]. Phages infect bacteria, killing them or becoming part of their genome [8]. Thus, phages are essential in shaping microbial communities in many ecosystems, including the human gut [9]. Phages are abundant in the human gut, ranging from 10^9^ to 10^12^ VLPs per gram of feces [10, 11, 6]. They can interact indirectly with the host through the modulation of bacterial gut communities or directly by adhering to mucins, inhibiting bacterial colonization and infiltration of the underlying tissues [12], and stimulating host immune activity. Changes in the diversity and richness of the gut phage populations have been linked to obesity and metabolic syndrome in children and adults [6, 7]. Additionally, dietary interventions, such as HFD, can impact the gut virome structure in humans and mice [13, 14].

There is a growing interest in transferring VLPs isolated from the fecal microbiota in a process called fecal virome transplantation (FVT). The FVT could alter gut bacterial richness and diversity in mice-to-mice assays [15, 16, 17, 18]. FVT has also shown promising results in treating metabolic diseases such as type 2 diabetes (T2D) and obesity, decreasing weight gain, and normalizing blood glucose tolerance in diet-induced obese mice [19]. It has also been demonstrated that FVT can change fecal microbiota and induce lean and obese body phenotypes in mice [20]. On the other hand, human studies have found that FVT from healthy and lean donors can improve glucose levels in individuals with MetS [21]. Furthermore, transferring VLPs from healthy donors can cure individuals infected with *C. difficile* [18]. However, these experiments only involved the FVT from mice to mice and human to human. A recent study found that transferring VLPs from human donors can alter gut-colonizing bacteria in a gnotobiotic mouse model with a humanized microbiota, modulating inflammatory bowel disease (IBD) symptoms [22].

This study examined the impact of administering fecal virus-like particles (VLPs) from normal-weight (NW) human donors to mice with obesity and MetS. The study focused on the impact of FVT on mice, their metabolic traits, and changes in their gut bacteriome and virome. To our knowledge, this is the first study to investigate the effects of FVT from human donors to mice with non-humanized gut microbiota to target obesity and MetS as proof of concept. This study opens the opportunity and provides valuable insights into using FVTs from humans in standard mice models to decipher meaningful bacteria-phage interactions associated with human diseases.

## Results

### 1. Inoculation of VLPs showed metabolic improvement in glucose tolerance in mice

First, we analyzed if using VLPs from normal-weight humans as inoculum for FVT could alleviate MetS symptoms in mice (Figure 1a). Twelve male C57BL/6 mice were fed a HFD for 14 weeks until they became obese and developed MetS. Then, mice were divided into two groups of 6 individuals each for the FVT treatment. The first group received a single oral dose of 4.4×10^8^ pooled fecal isolated VLPs (VLPs-inoculated group) obtained from 11 normal-weight human individuals (Figure 1a) [6]. The second group received only PBS buffer as inoculum (control group). Both groups continued with an HFD for 17 weeks (Figure 1b). After that, we found no significant difference in weight gain between the control and VLPs-inoculated groups (Figure. S1).

**Figure 1.**
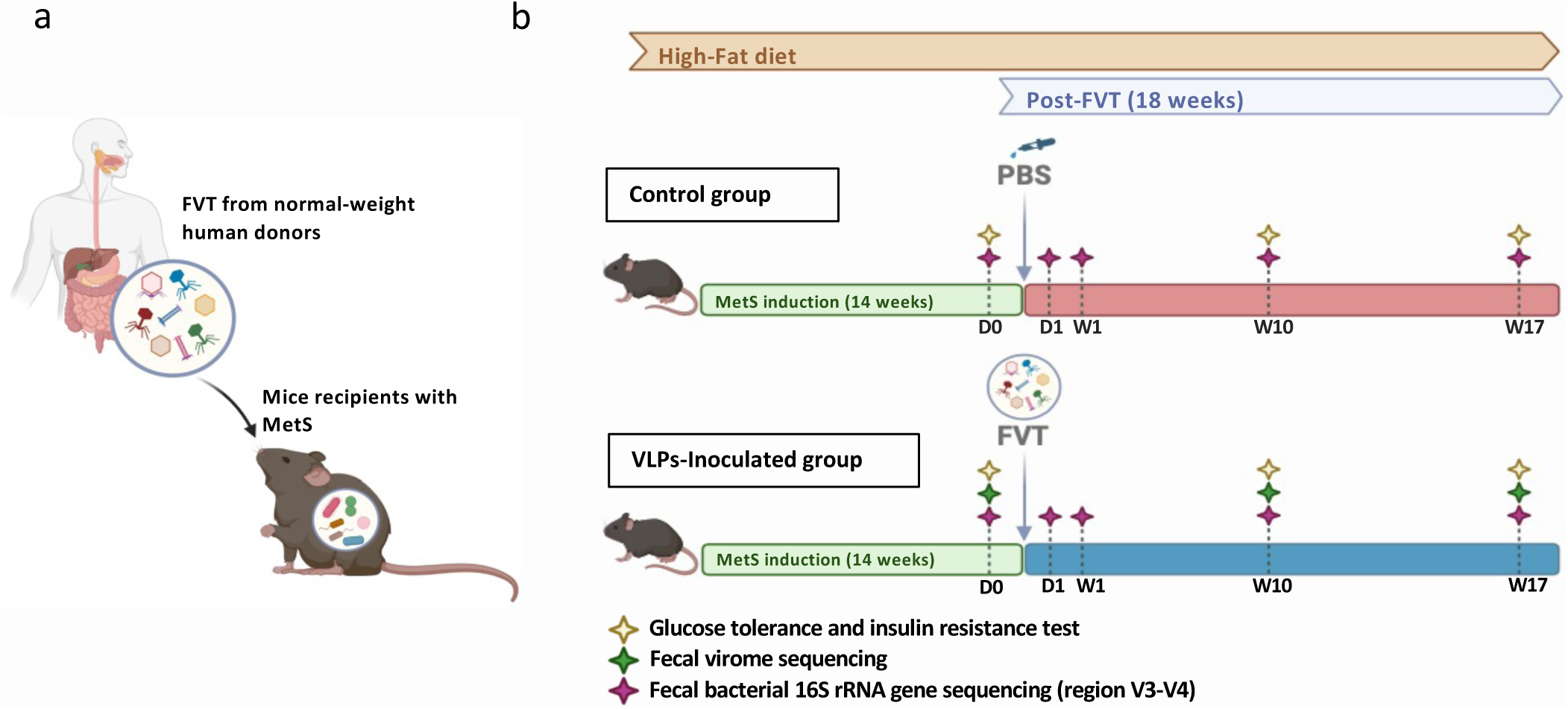
Overall description of the cross-infection model used in this study. a) The FVT was conducted by transferring the VLPs from normal-weight individuals to mice with unmodified gut microbiota. b) Experimental model for FVT in mice. The experiment involves twelve male C57BL/6 mice that were fed a high-fat diet (HFD) until they developed metabolic syndrome (MetS). The mice were then divided into two groups: the first group was given PBS orally as a control. At the same time, the second group was orally inoculated with VLPs isolated from fecal samples of normal-weight individuals (VLPs-inoculated group). The HFD was provided until the end of the experiment (17 weeks), and the weight of the mice was continuously monitored. Glucose tolerance and insulin resistance were tested pre-FVT on days 0 (D0) and post-FVT at weeks 10 (W10) and 17 (W17). The fecal bacterial microbiota was analyzed pre-FVT on day 0 (D0) and post-FVT on day 1 (D1) and weeks 1 (W1), 10 (W10) and 17 (W17). Additionally, the fecal virome was analyzed pre-FVT on days 0 (D0) and post-FVT at weeks 10 (W10) and 17 (W17).

We performed glucose tolerance tests (GTT) before (day 0) and post-FVT (weeks 10 and 17) to evaluate the glucose intolerance of the mice and observed a significant decrease in the area under the curve (AUC) of glucose concentration at week 17 compared to day 0 (Fig. 2a). This suggests that only the VLPs-inoculated group significantly improved blood sugar regulation. Moreover, the blood glucose levels were significantly lower at week 17 compared to day 0 and week 10 for the VLPs-inoculated group 60 minutes after glucose intake (Fig. 2b). Interestingly, we also observed a notably flattening effect of the glucose curve at week 17 (Fig. 2b). These findings suggest a significant improvement in glucose tolerance after 17 weeks post-FVT, while the control group did not show differences throughout the experiment, maintaining the curves for the three times very similar (Fig. 2c). Finally, we did not find significant differences among treatments in the insulin-resistance test (Fig. S2).

**Figure 2.**
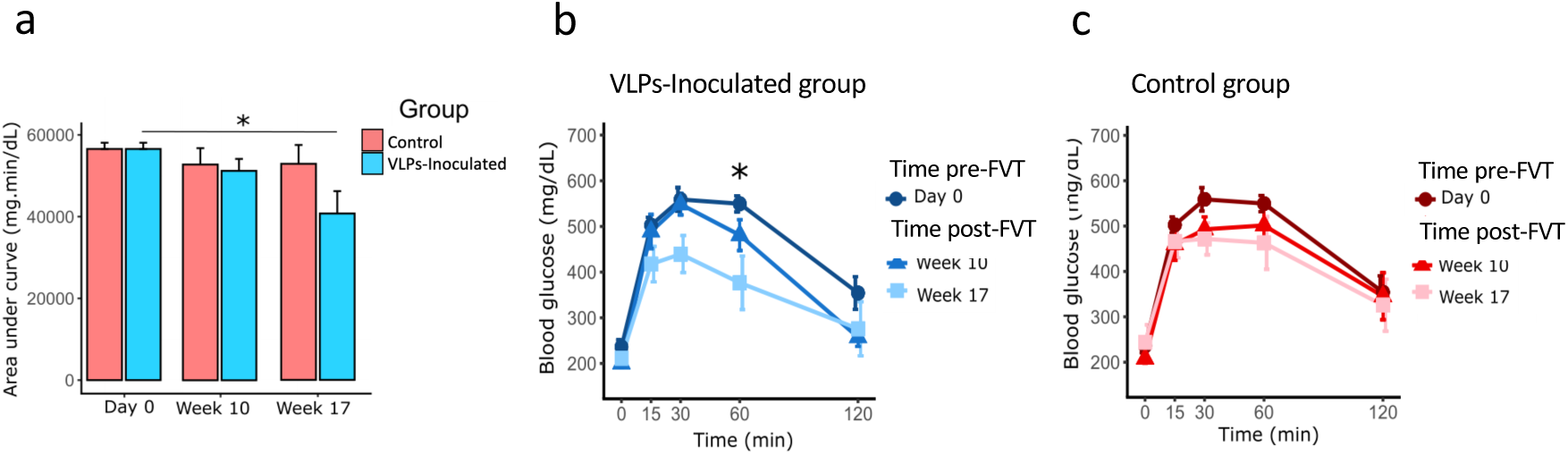
Results of the Glucose Tolerance Test. In part a), the graph displays the median of the area under the curve (AUC) of glucose concentration at different time points. In parts b and c, the graphs show the mean blood glucose concentration curves before (0 min) and after (15, 30, 60, and 120 min) intraperitoneal glucose administration to two groups of mice (N=6 each group). b) The VLPs inoculated group and c) the control group. The significance of the results is indicated by * p < 0.05.

### 2. VLPs transplant changed the overall gut microbiota and virome in mice

We examined the gut bacteria of mice by sequencing the V3-V4 regions of the 16s rRNA gene at day 0 and weeks 1, 10, and 17 post-FVT (Fig. 1b). After removing poor-quality reads, we assigned an average of 50,370 reads per sample to 141 OTUs with a frequency of at least 0.1% (Table S1). The richness rarefaction curves showed that all samples approached the saturation plateau (Fig. S3), and the Good’s coverage estimation showed that more than 99.9% of taxa were identified in all samples, indicating that our sequencing data successfully recovered most bacterial communities.

At the beginning (day 0) and the end (week 17) of the experiment, both the VLPs-inoculated and the control groups showed similar levels of bacterial richness and diversity (Fig. 3 a,b). However, at week 1, we noticed a significant decrease in bacterial diversity and richness in the control group compared to the VLPs-inoculated group (Fig. 3a,b). However, this decrease was restored at week 10 and was sustained until week 17 for both groups.

**Figure 3.**
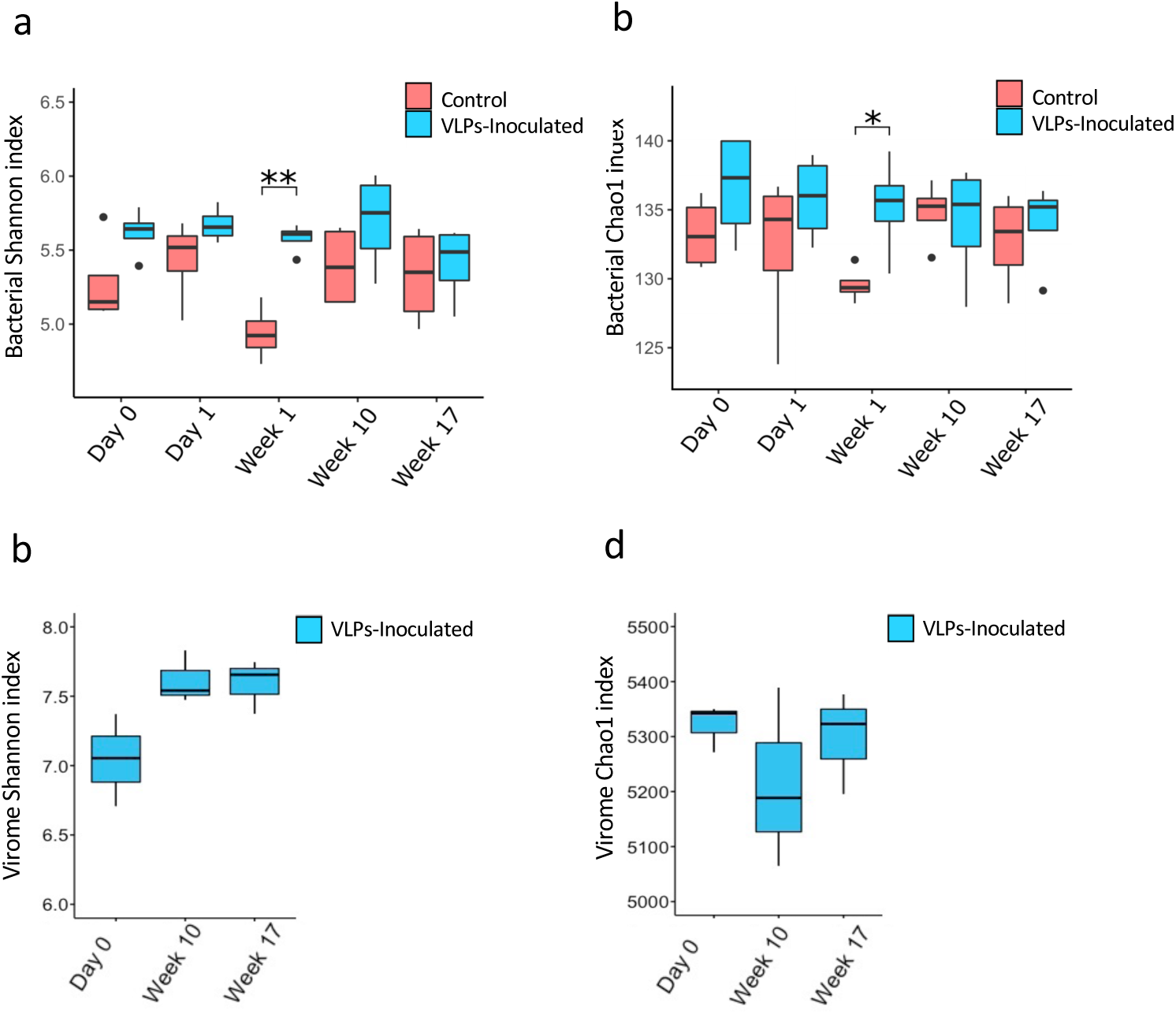
The alpha-diversity metrics. a) Shannon index and b) chao1 index of the microbiota for each group. Significance: * p < 0.05, ** p < 0.005. c) Shannon index and d) chao1 index of the virome for the VLPs inoculated group. Wilcoxon test did not observe a significant difference.

We also sequenced the fecal VLPs from the VLPs-inoculated group at days 0 and weeks 10 and 17 pos-FVT to explore the changes caused by the FVT in the original virome mice. After quality filters, we obtained an average of 2,701,536 paired-end sequences per sample. To ensure we only considered sequences potentially derived from VLPs, we discharged reads mapped to bacterial (∼29.63%) and mice (∼10.61%) genomes for further analyses and obtaining 14,173,277 total reads, averaging 1,574,808 reads per sample.

Our analysis revealed that phage diversity increased at week 10 and remained similar at week 17 post-FVT. However, it was not significantly different from day 0 (p = 0.076 and 0.084, respectively) (Fig. 3c). Moreover, the initial viral richness (day 0) was not significantly affected at weeks 10 and 17 post-FVT (Fig. 3d).

The beta diversity analysis of bacterial composition using UniFrac unweighted measures considering all samples revealed significant differences in microbial community compositions between the Control (Cluster 1) and VLPs-inoculated (Cluster 2) groups (Fig. 4a). The ANOSIM analysis confirmed a significant difference between the samples of two clusters (R= 0.325, p = 0.001).

**Figure 4.**
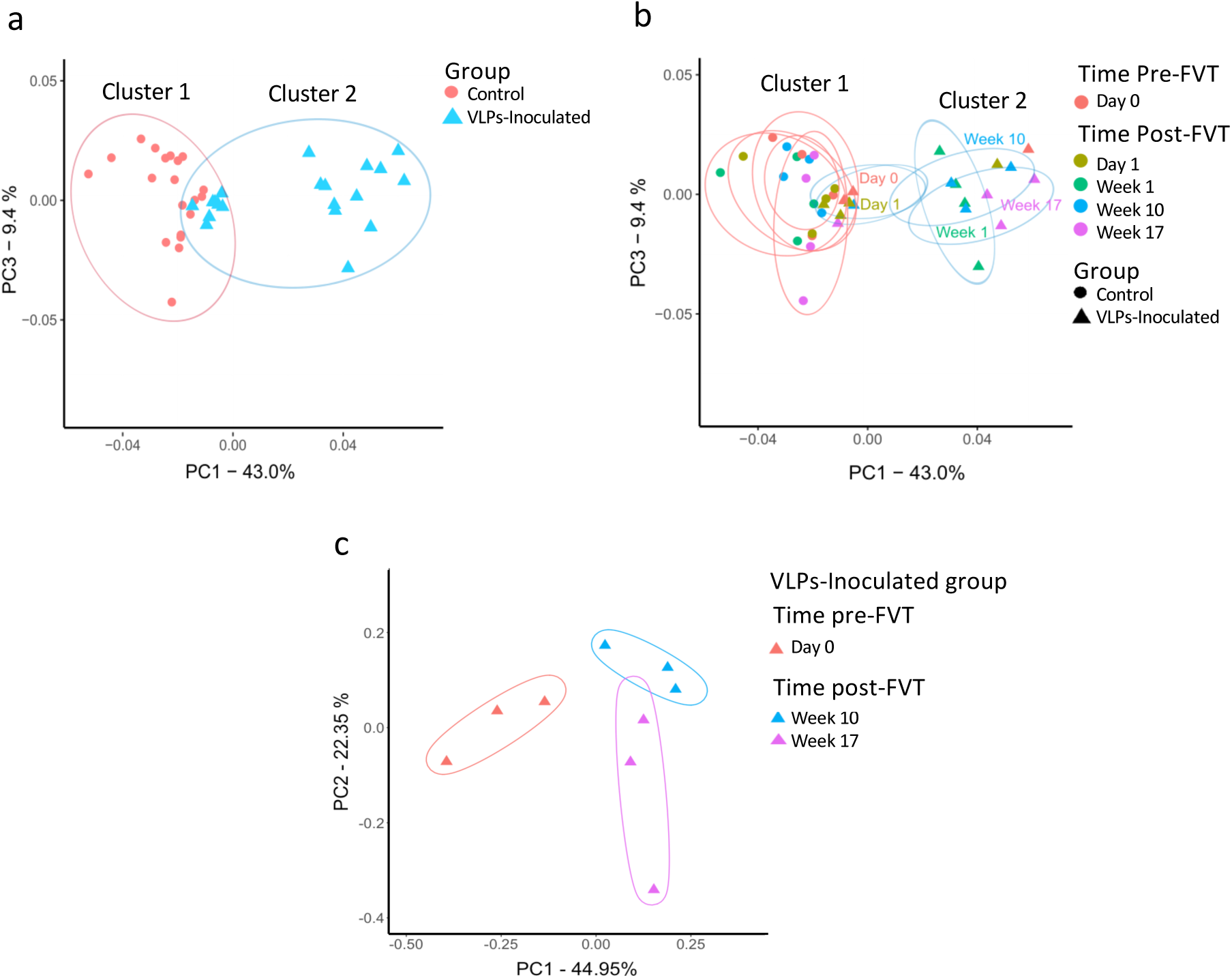
Beta diversity analysis for microbiota (a and b) and virome (c). The Unweighted UniFrac distances were used for all Principal Coordinate Analysis (PCoA) plots. a) Bacteria samples tagged by groups. b) Bacteria Samples were tagged by times pre- and post-FVT and by groups. c) Samples of the virome from the VLPs-inoculated group at different times pre and post-FVT.

It is worth noting that cluster 1 contained all the samples of the control group (from day 0 to week 17), suggesting no significant changes in the mice microbiota during the experiment for this group (Fig. 4a and 4b). Additionally, cluster 1 contained most of the samples of pre-FVT (day 0) and day 1 post--FVT from the VLPs-inoculated group (cluster 1, Fig. 4b). This suggests that on day 1, the FVT did not cause significant changes in the microbiota, being similar to the control group. On the contrary, cluster 2 contained no control samples, and most of the samples from weeks 1, 10, and 17 post-FVT from the VLPs-inoculated group (cluster 2, Fig. 4b). The ANOSIM confirmed a significant difference between the control and VLPs-inoculated groups at weeks 1, 10, and 17 (R= 0.432, p = 0.001). On the contrary, there was no significant difference between the groups on days 0 and 1 (R= 0.198, p = 0.064). These results suggest that after week 1 post-FVT, the microbiota composition of the VLPs-inoculated group significantly changed, separating from the control group, an effect maintained until weeks 10 and 17 (Fig. 4b).

On the other hand, we observed that the viral community at weeks 10 and 17 post-FVT significantly differed from the initial viral profile observed on day 0 (Fig. 4c). The ANOSIM statistical test confirmed that the three groups (day 0, weeks 10, and 17 post-FVT) were significantly distinct from each other (ANOSIM, R = 0.679; p= 0.009). This drastic change in the virome profile coincides with the changes observed at weeks 10 and 17 pos FVT in the microbiota, which also significantly differed from the initial bacteriome profile (Fig. 3b).

### 3. Specific bacterial changes were associated with the VLPs transplant

We compared the bacterial abundance between the control group and the VLPs-inoculated group at different times post-FVT to explore the enrichment of specific bacteria at each time point. As suggested by the beta-diversity analysis, the results showed no significant differences in bacterial abundances on days 0 and 1 post-FVT. However, there were significant differences at weeks 1, 10, and 17 post-FVT between the two groups (Fig. 5). At week 1, *A. muciniphila* was decreased in the VLPs-inoculated compared to the control group (0.0035 vs. 7.13%, log2FC= - 10.9, FDR= 1 x 10-11). Interestingly, this bacterium was also significantly decreased at weeks 10 (0.013 vs. 4.9%, log2FC=-8.24, FDR=6×10-5) and 17 (0.001 vs. 5.0%, log2FC =-11.9, FDR= 8×10-10) post-FVT (Fig. 5). A similar significant decrease in abundances was also observed for the highest taxonomic levels of this species, such as Akkermansia, Verrucromicrobiaceae, Verrucomicrobiales, Verrucomicrobiae, and Verrucomicrobia in the VLPs-inoculated group (Fig. 5).

**Figure 5.**
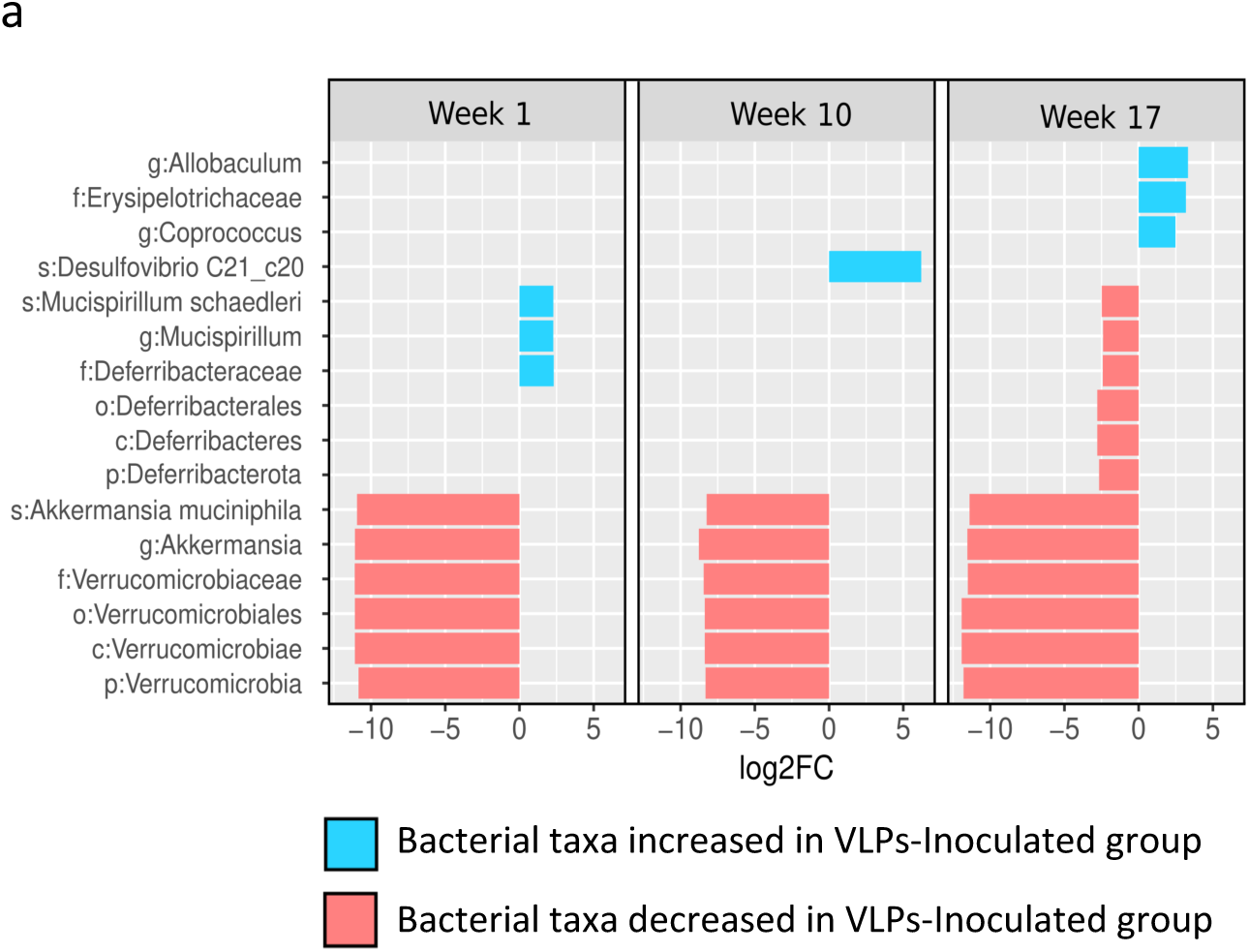
Analysis of the differential abundance of the microbiota at different times post-FVT. The blue bars indicate the significantly increased taxa, while the red bars indicate a decrease in the VLPs-inoculated group compared to the control group. The DESeq2 method was used to identify only those taxa with FDR<0.05 and a Log2fold change (Log2FC) cut-off of 1.5. The abbreviations used in this analysis are p: phylum, c: class, o: order, f: family, g: genus, s: species. On day 1 post-FVT, no significant differences in bacterial abundance between the two groups were observed (data not shown).

Also, the VLPs-inoculated group had a significantly increased abundance of *Mucispirillum schaedleri* (phylum Deferribacteres) at week 1 post-FVT, compared to the control group (1.1% vs. 0.15%, logFC=2.28, FDR=0.008) (Fig. 5). However, this abundance significantly decreased at week 17 post-FVT (0.33% vs. 2%, log2FC=-2.48, FDR=0.02). Furthermore, we observed a significant increase in Desulfovibrio C21_c20 at week 10 post-FVT, while Allobaculum, Erysipelotrichaceae, and Coprococcus were also significantly increased at week 17 post-FVT in the VLPs-inoculated group (Fig. 5). Although, it is worth noting that the abundance of Coprococcus progressively increased in the VLPs-inoculated group from day 1 to week 17 post-FVT (Fig.S4).

### 4. Specific virome changes were associated with the FVT

Out of the 5,498 viral contigs assembled with a length greater than 2kb, 2,222 were identified as potential phages at the family level (Fig. 6a). Most phage contigs were classified as Siphoviridae, Myoviridae, and Caudovirales, regardless of the experimental time point. Although there were slight differences in family abundances throughout time, we did not find any significant differences between pre-FVT (day 0) and weeks 10 and 17 post-FVT in the VLPs inoculated group (Fig. 6b). However, when we analyzed the abundance of each viral contig independently of their classification, 1,203 phage contigs showed significant differences between week 10 (Fig. 6c) and week 17 post-FVT (Fig. 6d) compared to day 0. Of these, 441 and 237 viral contigs were significantly enriched at week 10 and 17 post-FVT, respectively, and 142 viral contigs were commonly overabundant in both time points (Fig. S5). The unsupervised hierarchical clustering also confirmed the distinct virome profiles grouping the samples according to the time post-FVT (Fig. 6c, d).

**Figure 6.**
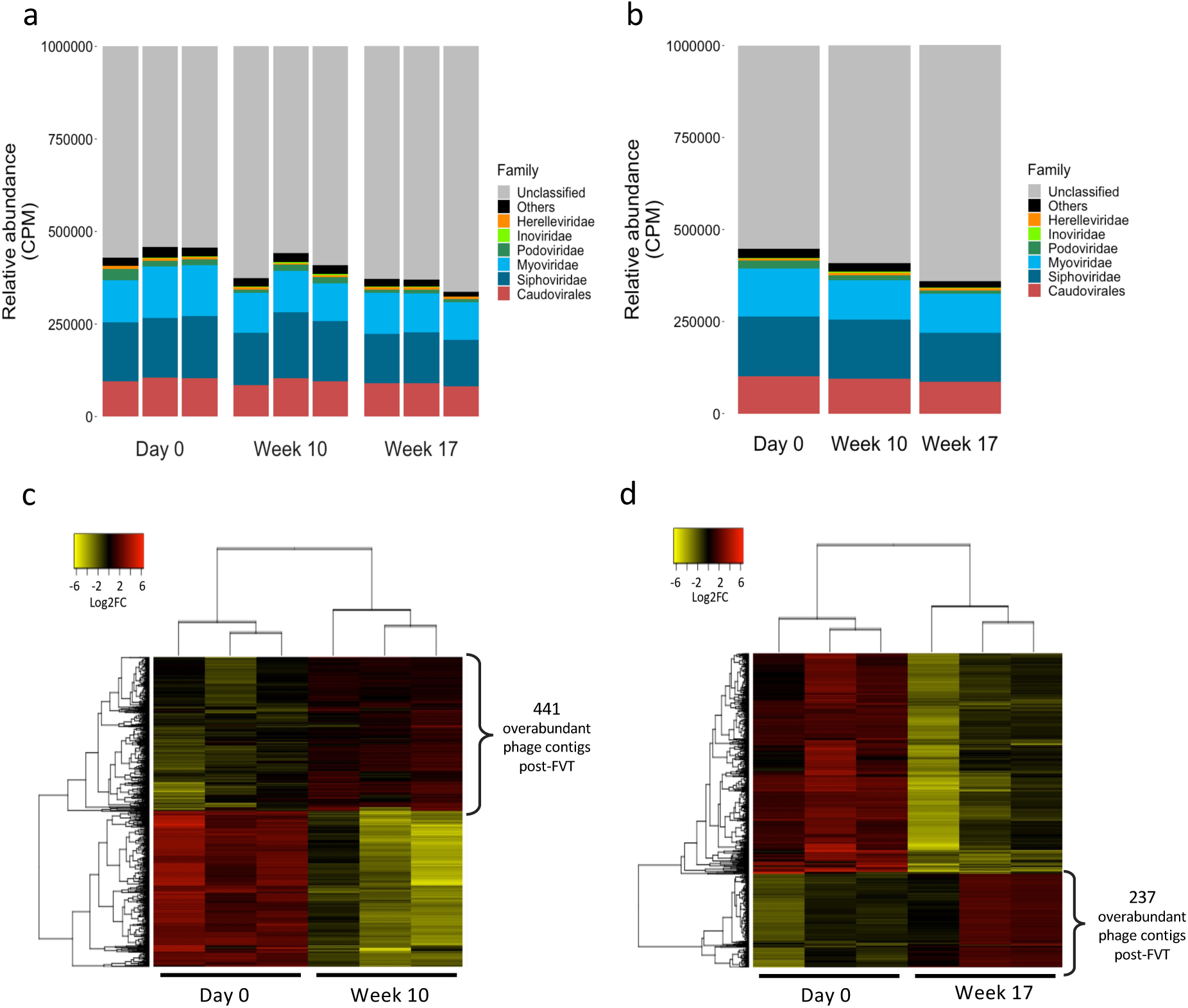
Summary of the viral classification of the virome and the overabundant phage contigs. Barplot of relative abundance normalized to counts per million (CPM) of viral families in VLPs-inoculated group by sample in a) and b) the average for each time point of the experiment. Unsupervised hierarchical clustering analysis of phage contigs abundance for the 1,203 differentially abundant between VLPs-inoculated and control groups. The heatmap shows the CPM abundances of individual phage contigs across the samples. c) differentially abundant phage contigs comparing week 10 against day 0, and d) differentially abundant phage contigs comparing week 17 against day 0. Heatmap colors represent CPM abundance, as indicated in the color key. The overabundant phage contigs of week 10 and week 17, as compared to day 0, are signaled in bullets.

### 5. The abundances of MetS biomarkers correlated with differentially abundant bacteria

We explored whether changes in bacteria due to VLP transplant were linked to changes in weight, GTT, and insulin resistance test measurements. Out of the 16 differentially abundant taxa (Fig. 5a), the abundance of A. muciniphila through experiment (Fig. 7a) showed a negative correlation with weight (r=-0.36, p=0.04) (Fig. 7b). In contrast, the corresponding family, order, class, and phylum also showed similar correlations. In contrast, a positive correlation was observed between the abundance of A. muciniphila and AUC glucose values of GTT (r=0.46. p=0.04) (Fig. 7C). Interestingly, after one week post-FVT, the abundance of A. muciniphila significantly decreased and was absent in all samples at week 17 post-FVT (Fig. 7a).

**Figure 7.**
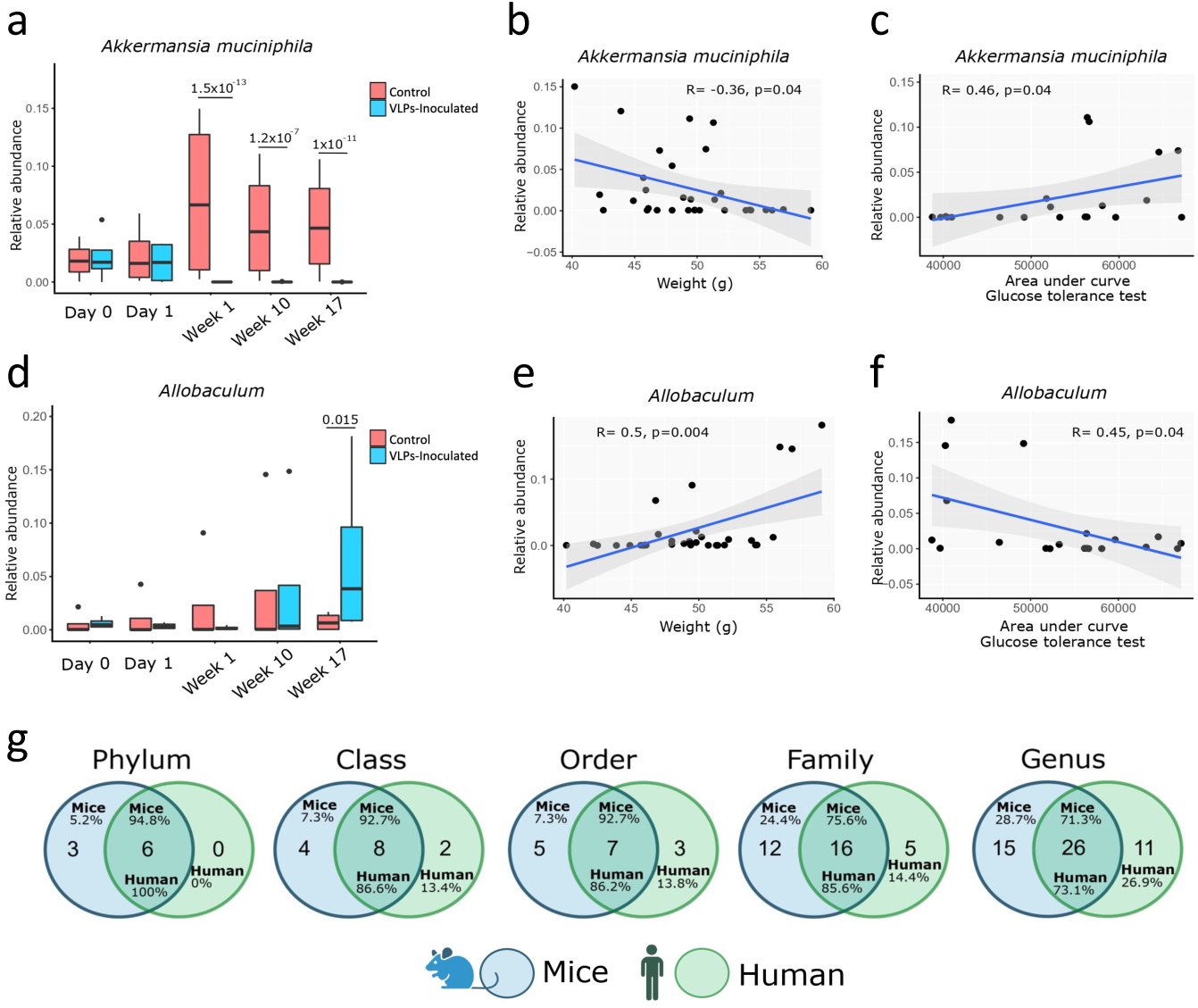
Abundances and correlations of critical bacteria enriched and depleted pos-FVT and microbiota shared between human donors of VLPs and model mice. a) The relative abundance of *Akkermansia muciniphila* across time points of the experiment pre and post-FVT in VLPs-inoculated and control groups. Spearman correlation of *Akkermansia muciniphila* and b) weight and c) area under the curve (AUC) of glucose levels in mice. d) The relative abundance of Allobaculum across time points of the experiment pre and post-FVT in VLPs-inoculated and control groups. Spearman correlation of Allobaculum and e) weight and f) area under the curve (AUC) of glucose levels in mice. g) Venn diagrams comparing bacterial taxa shared between human donors of VLPs and mice used in this study. The numbers inside the circles indicate the count of bacterial taxa unique to either humans (in blue) or mice (in green). The intersection of the circles represents shared taxa between groups, and the percentages indicate the abundance of shared taxa in each corresponding data set. Only the taxa with a relative abundance greater than 0.01% were considered.

On the contrary, the abundance of Allobaculum through experiment (Fig. 7d) had a positive correlation with weight (r=0.5, p=0.004, Fig. 7e). This correlation was observed not just at the genus level but also at higher taxonomic levels such as Erysipelotrichaceae, Erysipelotrichales, and Erysipelotrichi. Furthermore, there was a negative correlation between Allobaculum and AUC glucose values of GTT (r=-0.43, p=0.04, Fig. 7f). It is worth noting that the abundance of Allobaculum progressively increased in the VLPs-inoculated group over weeks 10 and 17 post-FVT (Fig. 7d).

### 6. The overabundant phages correlated with the differentially abundant bacteria

To explore possible relationships between the gut bacteriome and the virome, we conducted a Spearman correlation analysis between the abundances of the 16 differentially abundant taxa (Fig. 5) and the 142 phage contigs that were commonly overabundant in week 10 and 17 post-FVT (Fig. 6 c,d and Fig. S5). We only considered correlations with a Spearman coefficient (ρ) ≥ 0.8 and an asymptotic p-value ≤ 0.05. Our analysis revealed associations between 13 contigs identified as phages and bacteria (Fig. S5).

### 7. The microbiota from VLPs human donors was similar to the mice recipients

When investigating the effect of phages derived from human microbiota on mice, it is essential to consider the similarity between the microbiotas of both species. This is because phages from human microbiota have a higher chance of attaching to host bacteria in mice if their microbiotas are similar. Thus, we examined the microbiota of human fecal samples from VLPs donors and the microbiota of the mice used for the FVT. Interestingly, we found a high percentage of overlapped taxa between both microbiotas (Fig. 7g). Notably, the shared taxa (intersection of Venn diagram) have the highest abundance in their respective microbiotas (Fig. 7g). For instance, at the genus level, the 26 genera common to both microbiotas accounted for 71.3% of the abundance in mice and 73.1% in human samples. These results suggest that phages from human microbiota used for the FVT could potentially have a bacterial host in the mice microbiota.

## Discussion

We conducted a pilot study to explore the effect of fecal VLPs transplantation from healthy humans on mice with MetS. Although we found no significant difference in weight gain between the VLP-inoculated and control groups, the FVT significantly improved mice glucose tolerance despite being maintained on HFD. Both groups showed similar weight gain during the experiment, which could be attributed to the prolonged HFD (14 weeks pre-FVT and 17 weeks post-FVT). Recent studies have shown that FVT can drive lean and obese body phenotypes in mice, leading to weight gain or loss [20]. Additionally, Rasmussen et al. found that VLPs from normal-weight mice donors transplanted into HFD mice significantly decreased weight gain [15]. However, there were significant differences between their study and ours. Specifically, they used mice as FVT donors, while we used humans. Their mice were fed an HFD six weeks before the FVT, while ours were fed for 14 weeks. The number of VLPs administered in their study was 100 times greater than ours. Lastly, Rasmussen used two VLP inoculations in the FVT, while we used a single initial dose. All these differences could be associated with the lack of impact on mice weight observed in our study. We observed an improvement in glucose tolerance in the VLPs-inoculated group starting from week 6 post-FVT and being significant at week 17 compared to Day 0 (Fig. 2), with no changes in insulin resistance despite maintaining the HFD. Rasmussen also observed a substantial improvement in glucose tolerance after six weeks of FVT. It is worth noting that previous studies show FVT in individuals with obesity, and MetS does not decrease their body weight [15].

The mice used in our study were fed an HFD diet for 14 weeks until obesity and MetS developed. After that, their microbiota was characterized by a higher proportion of Firmicutes, which correlated with the animals’ weight gain, and a lower proportion of Bacteroidetes, which showed an inverse correlation with weight. The VLPs-inoculated group showed a significant change in the microbiota profile in the first week post-FVT, which was not observed in the control group (Fig 4b). Earlier studies in gnotobiotic mice showed that microbiota changes could be triggered by the inoculation of VLPs [23] or by administering specific lytic bacteriophages [9]. These studies found significant changes in the microbiota within the first-week post-FVT, but they only investigated the microbiota dynamics up to 2-3 weeks post-FVT. Another study reported that the ileum microbiota of normal-weight mice inoculated with VLPs from obese mice tends to change towards a microbiota characteristic of obesity on the fourth-day post-FVT [17]. The studies mentioned above need to be evaluated for a more extended period post-FVT to determine if the observed changes in the microbiota are maintained. According to our study, the changes in the microbiota resulting from the VLP inoculation were still present up to 17 weeks post-FVT (Fig. 4b).

The beneficial or detrimental impact of A. muciniphila on the host can be associated with diet, showing that a prolonged HFD (7 months) can increase the abundance of this bacterium [24]. A. muciniphila was increased in our control group from week 1 to week 17 (Fig. 7a), suggesting that prolonged HFD could be associated with this increase. On the contrary, a significant decrease in A. muciniphila was observed in the VLPs-inoculated group over the time post-FVT (Fig. 7b), suggesting VLPs could reverse the adverse effects of the HFD via depletion of this bacterium. A diet lacking fiber over time causes an overabundance of A. muciniphila, which can grow by breaking down the mucins that create the intestinal mucosal barrier, causing harm to the intestinal epithelium [25]. The loss of intestinal epithelium was associated with the development of MetS [26]. This damage makes the body more susceptible to inflammatory responses and susceptibility to pathogens [27].

In our study, mice were fed an HFD for 14 weeks, followed by another 17 weeks post-FVT. The HFD contained 6% fiber (a low amount), which may have contributed to the harmful effects of the overabundance of A. muciniphila observed in the control group that maintained the MetS. Notably, the VLPs-inoculated group experienced a significant decrease in the abundance of A. muciniphila post-FVT (Fig. 7a), which may have contributed to the observed improvement in glucose tolerance (Fig. 2). In this regard, the abundance of A. muciniphila was positively correlated with the AUC of glucose in GTT (Fig. 7C), meaning this bacterium could be associated with glucose intolerance. Interestingly, it has been observed that A. muciniphila was over-abundant in patients with type 2 Diabetes Mellitus [28]. However, other studies have reported the opposite trend [29, 30]. An overabundance of A. muciniphila has been associated with ulcerative colitis [31], colorectal cancer [32], Parkinson’s [33], and multiple sclerosis [34].

The VLPs-inoculated group showed a significant increase of *Mucispirillum schaedleri* one week post-FVT (Fig. 5). It has been reported that an HFD increases the abundance of M. schaedleri, causing inflammation in mice [24]. Thus, the over-abundance of this bacterium in our VLPs-inoculated group could be associated with prolonged exposition to the HFD (14 weeks pre-FVT and 17 weeks post-FVT). However, during week 17 post-FVT, M. schaedleri experienced a significant decrease in the VLPs-inoculated group (Fig. 5). M. schaedleri was also early associated with proinflammatory activity [35]. Thus, we suggest that FVT helped to inhibit this activity in the VLPs-inoculated group and could be linked to the improvement in glucose tolerance of our mice model. The decrease in A. muciniphila and M. schaedleri in the VLPs-inoculated group could be associated with a lytic effect of the FVT on these bacteria. Alternatively, the abundance of other non-susceptible bacteria could have been affected by the depletion or increase in susceptible bacteria to lytic phages due to inter-bacterial interactions [9].

We also found that the Allobaculum significantly increased in the VLPs-inoculated group at week 17 (Fig. 5 and 7d). Allobaculum improves MetS in obese mice and rats when given prebiotics and probiotics [36, 37]. This beneficial effect could be related to their short-chain fatty acid production, such as butyrate, which is good for the host’s health and metabolism [38]. This bacterium also reduced proinflammatory markers such as IL-1β and TNF [37]. The increase of Allobaculum observed in our VLPs-inoculated group might compensate for the lack of A. muciniphila in producing short-chain fatty acids. While A. muciniphila is limited to using mucins as a sole energy source, Allobaculum can metabolize glucose to produce lactate or butyrate. In addition, Allobaculum is highly active in using glucose, which could contribute to their rapid catabolism [39]. Interestingly, this bacterium negatively correlated with the AUC of glucose in GTT (Fig. 7f), suggesting that Allobaculum could have a beneficial hypoglycemic effect on the mice inoculated with VLPs even though they were maintained in an HFD.

The group inoculated with VLPs had an overabundance of Coprococcus at all times post-FVT (Fig S4), being significantly overabundant at the end of the treatment (week 17) (Fig. 5 and Fig. S4). Studies have shown that an overabundance of Coprococcus can improve quality-of-life indicators in mental health, and it also was depleted in depression [40]. This bacterium has also been found to promote the desire to exercise in mice [41]. Additionally, Coprococcus has also been suggested as a probiotic for Inflammatory Bowel Disease (IBD) as it was significantly increased during IBD recovery [42]. Based on these results, it can be suggested that Coprococcus is beneficial in mitigating the MetS effects in our VLPs-inoculated group.

The human gut virome can vary significantly among individuals [6]. We used a mix of viruses from multiple healthy individuals to ensure a highly diverse virome as inoculum for FVT in mice. This inoculum changed the virome populations at weeks 10 and 17, being different from the originality found at the beginning of the experiment (Fig. 4c). However, to discard some fecal bacteria from the donors may have been carried over into the mix, impacting the changes observed in the gut microbiome, we tested the VLPs mix for cultivable bacteria and observed no growth was found (data not shown). The virome analysis showed that the richness decreased in week 10 post-FVT in the VLPs-inoculated group compared to the pre-FVT (day 0). However, it recovered during week 17 post-FVT to the values observed pre-FVT (Fig. 3d). The virome diversity showed a slight increase in week 10, which remained consistent until week 17 post-inoculation compared to the pre-FVT (Fig. 3c). However, both changes were insignificant.

The Siphoviridae family was the most common in the virome, followed by Myoviridae and Podoviridae (Fig. 6a,b). These families have been previously found in mice and human viromes. However, Podoviridae was more abundant in other mouse viromes [43]. Our samples found that Podoviridae was less abundant, which may be due to our use of a murine model of obesity. On the other hand, Herelleviridae and Inoviridae were found in low numbers in our samples (Fig. 6a,b), while they were not found in other mouse virome studies [43]. We did not detect significant changes in the abundance of viral families over time, but we observed that some particular phages were differentially abundant at weeks 10 and 17 (Fig. 6C and D). Although we did not observe significant changes in the abundance of viral families over time, we did notice that particular phages were over-abundant at weeks 10 and 17 (Fig 6c, d), suggesting a phage-bacteria dynamics over the time post-FVT.

The FVT has shown potential in treating various human diseases, such as C. difficile infections [18], and improving glucose levels in people with MetS [21]. In mice, the FVT reduced intestinal bacterial overgrowth [17] and improved weight and glucose tolerance in type 2 diabetes [15]. The FVT can also change gut bacteria and drive lean and obese body phenotypes [20]. However, these experiments only involved FVT from mice to mice and human to human. Recently, FVT from human donors was used to alleviate IBD symptoms by altering the gut-colonizing bacteria in the gnotobiotic mouse model [22]. This is the only study to date demonstrating the effectiveness of human VLPs in altering the gut microbiota of mice. However, the microbiota was previously humanized. Our study provides a proof of concept for the efficacy of FVT from human donors to improve glucose tolerance in mice by altering the gut microbiota and virome. Our study suggests that trans-kingdom interactions between the virome and bacteriome communities can be translated from human to mouse.

When mice received VLPs, we noticed a significant change in the recipient’s virome profile (Fig. 4c). This led us to believe that the viral community transferred from the donor could change the existing bacteriophages. We also discovered that phages derived from healthy individuals could interact with the mice bacteriome (Fig. S5). The microbiota from the VLPs donors was similar to that of the experimental mice (Fig. 7g), which could have contributed to the FVT engraftment in mice. Changes in the phage community of mice after FVT could be due to the establishment of donor-invading phages that successfully find and infect a host in the recipient microbial community or by activating recipient prophages.

This study showed that transferring VLPs from human feces to mice resulted in significant changes in the microbiota, the virome, and the metabolism of the animals. More research is needed to determine the molecular mechanisms underlying these changes, identify the specific viruses involved, and explore this approach’s potential biotechnological and clinical applications. The correlation analyses between the bacteriome and virome post-FVT showed significant associations between bacteria and viruses that need to be studied in depth (Fig. S5). This suggests that FVT may affect the trans-kingdom interactions between different components of the gut microbiome.

We need to address some limitations in our study. First, we could not look for genomic signatures of phage-bacteria interactions because we used 16S rRNA amplicon sequencing. Second, we used whole viromes, not individual viruses, which made it hard to find specific phages that caused the effect in mice. Fecal metabolites could have been carried over into the VLPs, which could have affected the observed phenotypes. However, the probability of metabolite carryover was reduced by using a recent VLP purification protocol [44].

## Materials and methods

### Subject details and sample collection of VLPS

The fecal samples of the donors for FVT were collected from a group of 11 normal-weight children aged 7-10 [6]. The samples were processed to isolate VLPs using the method previously described by our group [44]. Briefly, ∼250 mg of each fecal sample was resuspended in 40 mL of sterile buffer SM (100mM Sodium chloride, 10mM Magnesium sulfate, 50mM Tris-HCl, pH 7.5) and centrifuged at 4,700 x g for 30 min at room temperature and centrifuged at 4,700 x g for 30 min at room temperature to precipitate large debris. The VLP-containing supernatant was filtered through a 0.45 μm PES syringe filter to remove the bacterial component. The resulting filtrate was concentrated to 200 μL with an Amicon Ultra 15 filter unit, 100 KDa (Cat. UFC910024, Millipore, MA, USA). Next, 40 μL chloroform was added to the concentrate and incubated for 10 min at room temperature to degrade cell debris. The non-capsid-protected DNA was eliminated with 2.5 units per mL of DNase I following the manufacturer’s procedure (Cat 18047-019, Invitrogen, MA, USA). The treated VLPs were then concentrated with AMICON and recuperated in 200 μL SM buffer. The purity and VLP concentration were assessed using SYBR Green staining and epifluorescence microscopy in an Olympus FV1000 Multiphoton Confocal Microscope. Each field was quantified in triplicate using the free image processing software Fiji. Finally, the VLPs from all 11 samples were mixed and used to inoculate the mice in the study.

### Animal study design

We used twelve male C57BL/6 mice of 3-4 weeks of age that were randomly mixed and fed a standard diet (3.1 kcal/g= 18.6% protein, 44.2% carbohydrates, and 6.2% fat; Teklad Global, 2018) for four weeks to homogenize gut microbiota. Afterward, the mice were divided into two experimental groups, “Control” and “VLPs” (6 mice per group), each group divided into two cages of 3 mice each. Both groups were fed a high-fat diet (HFD) for 14 weeks. The HFD (5.24 kcal/g= 20% protein, 20% carbohydrate, and 60% fat; Research Diets cat. D12492) was used to induce obesity and metabolic syndrome according to the previously standardized model by Manzo et al. (submitted). This model is characterized by obesity, weight gain, insulin resistance, glucose intolerance, and hypertriglyceridemia.

### Fecal Virome Transplant

We administered 100 µL of 1.33% NaHCO3 orally to neutralize the gastric pH of the animals for FVT. After that, we gave the Control group 200 µL of saline buffer, while the VLP group received 200 µL of a suspension containing the VLPs (4.4 x 10^8^ VLPs) from the 11 normal-weight children. Both groups were then kept on an HFD for 17 weeks until they were sacrificed.

### Blood plasma analysis

Blood plasma analysis was conducted by performing glucose tolerance tests (GTT) at day 0 and weeks 10 and 17 post-FVT and insulin resistance tests at weeks 10 and 17 post-FVT. During the GTT, glucose was administered, and blood was collected after 0, 15, 30, 60, and 120 minutes to measure its levels. Similarly, during the insulin resistance test, insulin was administered (1U/kg; Humulin R., Eli Lilly), and glucose levels were determined in the same way at the same time intervals. Glucose or insulin administration was performed intraperitoneally, and blood was collected through a tail puncture for measuring glucose levels.

### Sample collection and extraction of bacterial DNA

We collected fecal samples at five experiment points: day 0 (pre-FVT) and post-FVT on day 1, week 1, 10, and 17. For 16S gene sequencing, four mice per group (2 per box) were selected. The samples were stored in RNA Later solution (Invitrogen) at -70 °C until use. DNA extraction was performed with the Quick-DNA Fecal/Soil Microbe kit (Zymo Research). Agarose gel electrophoresis and Qubit (Invitrogen, Cat. Q33231, CA, USA) determined the DNA integrity and concentration, respectively. Next, the V3–V4 hypervariable region of the 16S rRNA genes was amplified using the universal primers 338F (5′-ACTCCTACGGGAGGCAGCAG-3’) and 533R (5’-TTACCGCGGCTGCTGGCAC-3′). The resulting ∼500 bp PCR products were then purified using magnetic beads (Agencourt AMpure XP, Beckman Coulter) and barcoded according to the Illumina Sequencing Library Preparation user’s guide. Finally, each library’s concentration and size distribution were assessed with a Qubit fluorometer and an Agilent 2100 Bioanalyzer (Agilent Technologies, Santa Clara, CA, USA). The libraries were sequenced in an Illumina MiSeq platform with a 2 × 250 Paired-End format at the Sequencing Unit from the National Institute of Genomic Medicine, Mexico.

### Bioinformatic analysis of bacterial DNA

The raw sequences obtained underwent quality control, including removing primer sequences from each read and filtering sequences with a quality score >20 and length >350 nt to ensure complete V3-V4 regions of the 16S rRNA gene [48]. All quality-filtered sequences were joined and analyzed using QIIME (version 1.9). The sequences were clustered into operational taxonomic units (OTUs) based on 97% sequence similarity against the Green Genes (GG) sequence database version 13.8. The singletons were excluded from downstream analyses. The OTUs table was filtered by keeping those with >0.1% abundance to eliminate rare, transient, or potential sequencing errors [49]. Alpha diversity indices (Shannon, Chao1, and Observed OTUs) were calculated using 10,000 rarefactions at the depth of the smallest sample. Beta diversity was determined with the Bray Curtis, Jaccard, and UNIFRAC distances (weighted and unweighted), and principal coordinate analysis (PCoA) was used to visualize the proximity (similarity) or distance (dissimilarity) of the samples in the graphs of two dimensions. Differential abundance analyses of bacteria between groups were performed with DESeq2 using the Microbiome Analyst web server [41].

### Comparison of human and mouse fecal microbiota

To assess the degree of overlap between the mouse fecal microbiota and that of humans used as donors of VLPs in this study, we employed their previously reported 16s sequences for this cohort of children [6]. We then compared the bacterial taxonomic groups identified in the mice from this study with those in the human dataset. All sequences underwent consistent processing using the described bioinformatics pipeline to obtain bacterial taxonomic assignments. The presence or absence of bacterial taxa between humans and mice was visualized using Venn diagrams, considering those with relative abundance > 0.01%.

### Processing of fecal samples for viral DNA extraction and sequencing

To analyze the virome, we selected the fecal samples of the FVT group at three experiment points: day 0 (pre-FVT) and post-FVT on weeks 10, and 17. We selected six mice (2 per box) for each time point, and the stools were paired and mixed at 50 mg per mouse, resulting in three pools per time. The VLP isolation was conducted using the previously reported extraction protocol [6]. Following the previously reported protocol, the VLP numbers were determined using epifluorescence microscopy [44]. The DNA of VLPs was then extracted, following the manufacturer’s protocol for the QIAamp MinElute Virus Spin Kit (Cat. 57704, Qiagen, Hilden, Alemania). The resulting DNA for each sample was quantified with a Qubit fluorometer (Cat. Q32851, Thermo Fisher Scientific, CA, USA) and diluted with ribonuclease-free water to a concentration of 0.3 ng/μl. Independent sequencing libraries were then prepared from this DNA, following the Illumina Nextera XT DNA Library Preparation protocol (Cat. FC-131-1024, Illumina, CA, USA). For DNA tagmentation, five μl of DNA at 0.3 ng/μl were mixed with the tagmentation reaction mix. Next, the indexed oligos were added and amplified in the library for 12 cycles. Finally, each library was purified with 30 μl of AMPure XP beads (Cat. A63881, Beckman Coulter, CA, USA) to obtain ∼600 bp DNA fragments. All barcoded libraries were sequenced using the Illumina NextSeq500 platform in the 2x150 pair-end mode at the National Institute for Genomic Medicine, México Sequencing Unit Facility.

### Bioinformatic analysis of viral DNA

Adapters from raw reads (max. 4,873,44 6reads and min. 2,073,324) were identified and removed using CUTADAPT [50], while low complex and dereplicated reads were eliminated using PRINSEQ [51]. Low-quality bases (PHRED Q20) were trimmed using TRIMMOMATIC [52]. The reads from humans were detected by read mapping using SMALT against the Homo sapiens GRCh38.p13 reference genome GenBank GCA_000001405.28 (parameters -c 60). The reads for bacteria and archaea were detected using Kraken [53] with default parameters. Then, all reads mapped to those genomes were removed, and the remaining reads were named quality reads. The quality reads from all samples (14,173,277) were pooled for de novo assembly using IDBA-UD assembler [54]with a kmer length of 35-115 and SPADES assembler with a kmer length of 35-115 with scaffolding rounds. Both assemblies were processed with METASSEMBLER, resulting in a meta-assembly. Scaffolds less than 2 kb were removed. Each sample’s reads were mapped separately with SMALT against the viral meta-assembly using -c 0.7, -y 0.9 parameters. Viral scaffolds covered ≥ 70% in length with 90 % of identity by the reads by at least one sample were used as a cutoff to discard chimeras. We used CD-HIT with a 90% clustering identity to eliminate the redundant contigs. The taxonomy classification of each contig was obtained using BLASTx against the NR NCBI viral protein database with a maximum e-value cutoff of 0.001 [45]. The taxonomy of each contig was assigned by the lowest-common-ancestor (LCA) algorithm in MetaGenomeANalyzer (MEGAN6), using the BLASTX results with the following parameters: Min Support: 1, Min Score: 40.0, Max Expected: 0.01, Top Percent: 10.0, Min-Complexity filter: 0.44. Scaffolds identified as eukaryotic viruses were removed from our analysis. Each sample’s reads were mapped with SMALT against the viral scaffolds using -c 0.7, -y 0.9 parameters. Raw counts matrix was input into DESeq2 to analyze differences before and after VLP gavage; scaffolds with adjusted p-value < 0.05 and a log2 fold change > 2 or < -2 were considered differentially abundant. QIIME performed bray-Curtis dissimilarity for diversity analyses, and analysis of similarities (ANOSIM) was used to evaluate multiple group comparisons. The alpha diversity Shannon and Chao1 index was determined, with R performing 10000 iterations on the raw counts matrix. The relative abundance of phage families was calculated on a matrix normalized by Counts per million (CPM), and linear discriminant analysis (LDA) among viral families was assessed with LEFSE.

### Statistical analysis

The students’ t-test was used to compare weight, glucose tolerance, and insulin resistance measures between mice groups. The area under the glucose curve (AUC) was calculated using the trapezoid method [46] with the DescTools R library. For comparisons in the beta diversity of the microbiota, the non-parametric ANOSIM method was used, employing 999 permutations, with a prior evaluation of the dispersion using Betadisper from the Vegan R library. The differences in the abundance of taxonomic groups were carried out by the DESeq2 method, taking as significant FDR values <0.05 and a log2fold changes (log2FC) cut-off point of 1.5 [47].

## Acknowledgements

We thank Juan Manuel Hurtado Ramírez for informatics technical support and the computational server maintenance, as well as Filiberto Sanchez for experimental technical support. M.C.E., thanks to the Doctoral Biochemical Sciences Program at IBT UNAM and CONACyT by doctoral fellowships C.V.U.: 887285. This work was supported by the CONACyT grant Ciencia de Frontera 2019 263986 and by the DGAPA PAPIIT UNAM (IN219723) to A.O.L. We would like to thank the postdoctoral fellowship to Fernanda Cornejo-Granados (CVU 443238) as part of the Estancias posdoctorales por México 2022 program. We also thank the donatives from DGAPA PAPIIT UNAM, IN213119 and IN217822 to L. P.M.; and IN211719 and IN216922 to G. P.A.

## Data availability

The data used in this study have been deposited under BioProject PRJNA1003883 to the NCBI database.

## Study approval statement

All animal experiments described in this study were approved by the Institutional Bioethical Committee, of Biotechnology Institute from National University Autonomous of Mexico; protocol number 300.

