## Supplementary information for "Fecal Virome Transplantation (FVT) from healthy humans remodels the gut bacteriome and virome and reduces metabolic syndrome in mice"

a

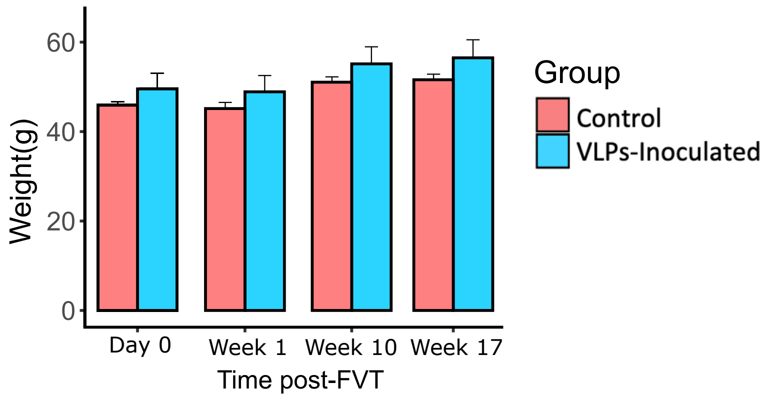


**Figure S1.** Weight of mice (g) with MetS pre and post-FVT of VLPs-inoculated and control groups. No significant differences were observed between groups.

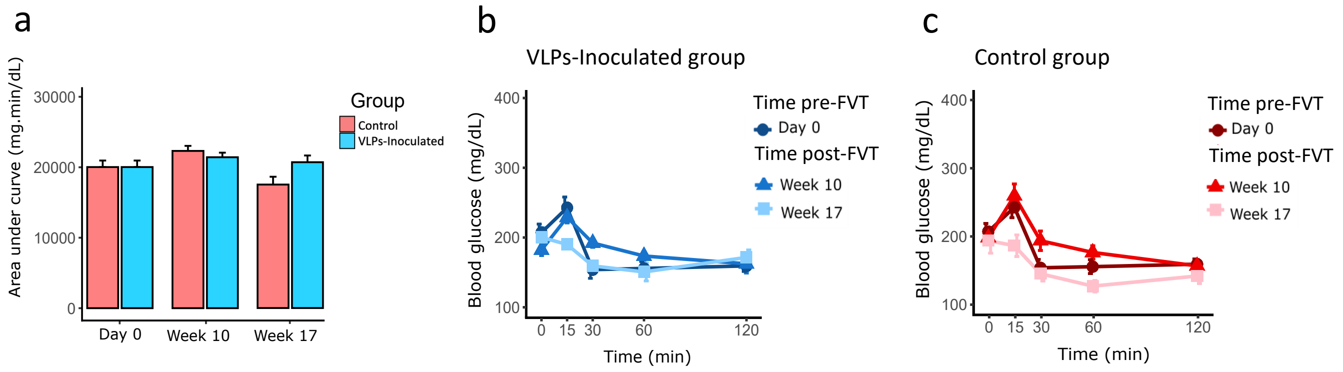


**Figure S2.** Results of the Insulin Resistance Test. In part a), the graph displays the median of the area under the curve (AUC) of glucose concentration at different time points. In parts b and c, the graphs show the mean blood glucose concentration curves before (0 min) and after (15, 30, 60, and 120 min) intraperitoneal glucose administration to two groups of mice (N=6 each group). b) The VLPs inoculated group and c) the control group. The significance of the results is indicated by * $p < 0.05$.

a

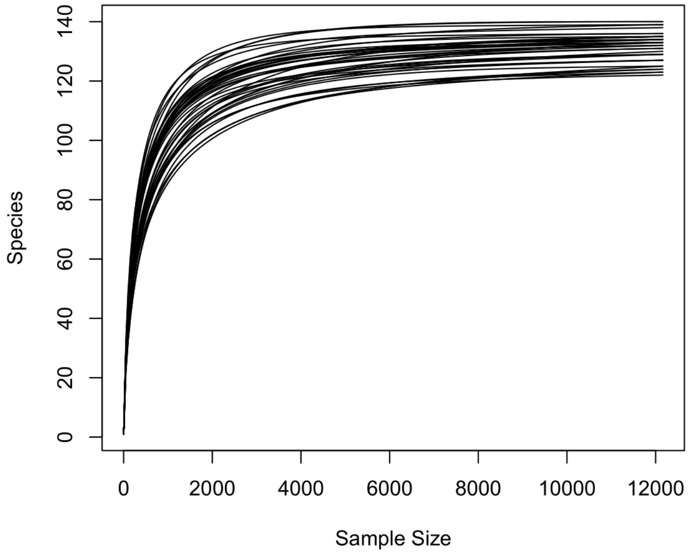


**Figure S3.** Alpha diversity rarefaction.

a

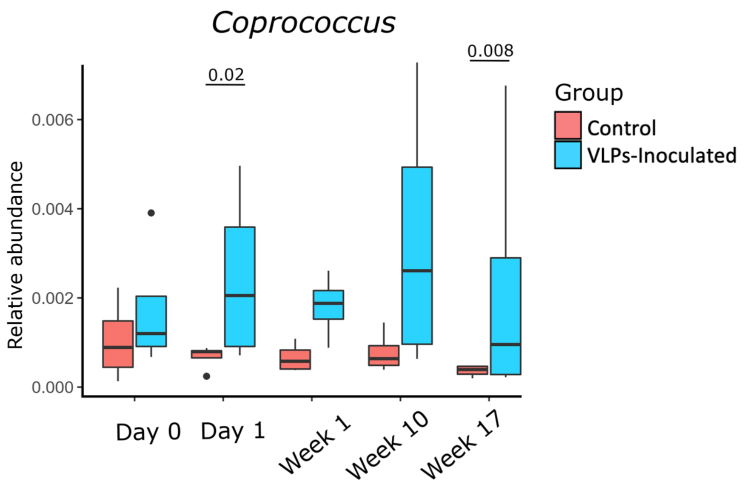


**Figure S4.** Relative abundance of the genus Coprococcus across time points of the experiment pre and post-FVT in VLPs-inoculated and control groups.

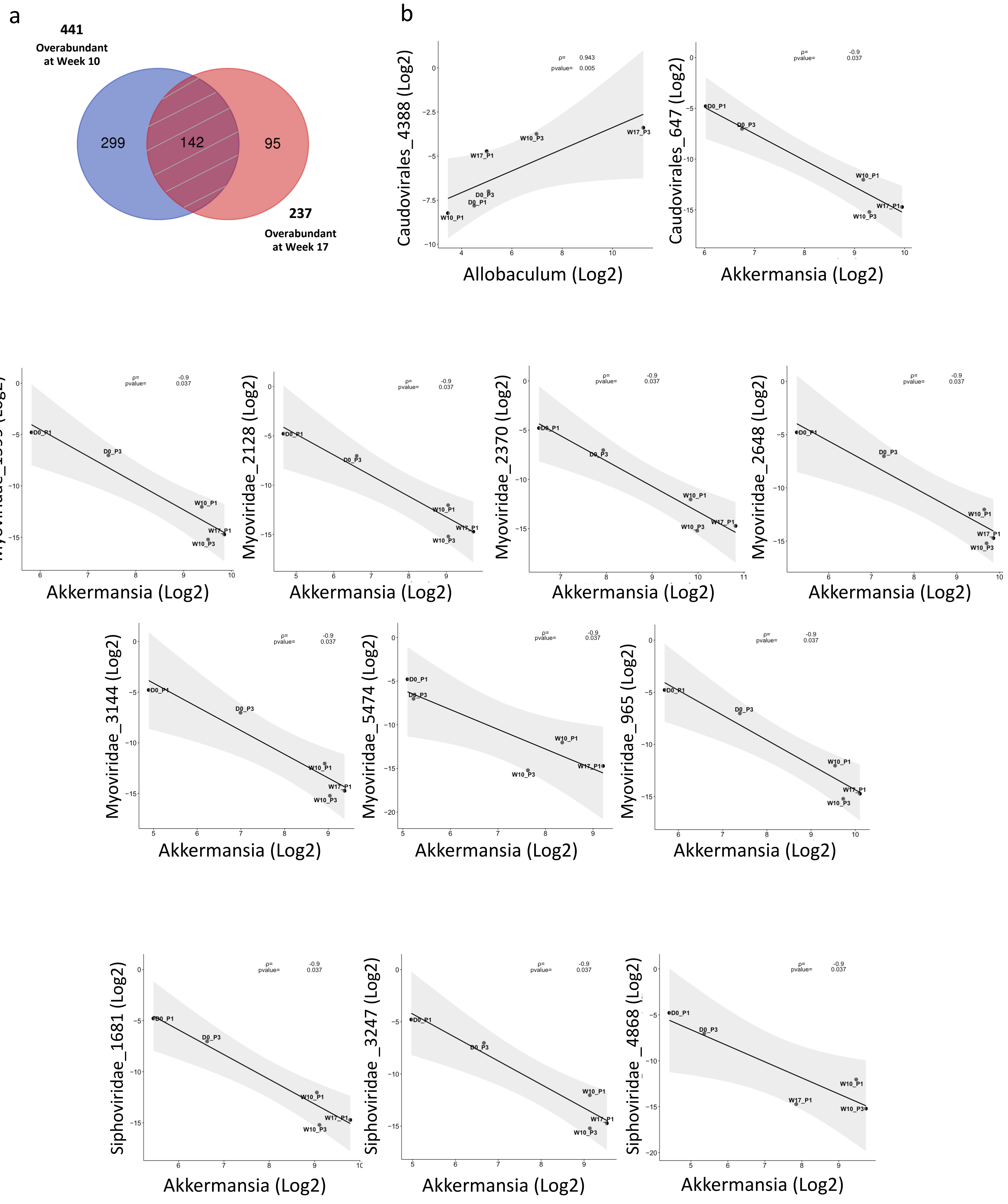
a
441
Overabundant at Week 10
299
142
95
237
Overabundant at Week 17
b
Caudovirales_4388 (Log2)
Allobaculum (Log2)
ρ= 0.943
pvalue= 0.005
Caudovirales_647 (Log2)
Akkermansia (Log2)
ρ= -0.9
pvalue= 0.037
Myoviridae_1599 (Log2)
Myoviridae_2128 (Log2)
Myoviridae_2370 (Log2)
Myoviridae_2648 (Log2)
Myoviridae_3144 (Log2)
Myoviridae_5474 (Log2)
Myoviridae_965 (Log2)
Siphoviridae_1681 (Log2)
Siphoviridae_3247 (Log2)
Siphoviridae_4868 (Log2)
D0_P1
D0_P3
W10_P1
W10_P3
W17_P1
W17_P3

**Figure S5.** a) The Venn diagram shows the 142 overabundant phage contigs shared in weeks 10 and 17 compared to day 0. b) Spearman correlation plots between the 142 overabundant phage contigs and the abundance of the differentially overabundant bacteria at different time points of the experiment.
